# Learning from All Views: A Multiview Contrastive Framework for Metabolite Annotation

**DOI:** 10.1101/2025.11.12.688047

**Authors:** Yan Zhou Chen, Soha Hassoun

## Abstract

Metabolomics, enabled by high-throughput mass spectrometry, promises to advance our understanding of cellular biochemistry and guide new discoveries in disease mechanisms, drug development, and personalized medicine. However, as the assignment of molecular structures to measured spectra is challenging, annotation rates remain low and hinder potential advancements. We present MultiView Projection (MVP), a novel framework for learning a joint embedding space between molecules and spectra by leveraging multiple data views: molecular graphs, molecular fingerprints, spectra, and consensus spectra. MVP builds on contrastive multiview learning to capture mutual information across views, leading to more robust and generalizable representations for spectral annotation. Unlike prior approaches that consider multiple views via concatenation or as targets of auxiliary tasks, MVP learns from all views jointly, resulting in improved molecular candidate ranking. Notably, MVP supports annotation using either individual spectra or consensus spectra, enabling flexible use of multiple measurements. On the MassSpecGym benchmark, we show that annotation using query consensus spectra significantly outperforms rank aggregation strategies based on constituent spectrum annotation. Using the consensus spectrum view, MVP achieves 35.99% and 13.96% rank@1 when retrieving candidates by mass and formula, respectively. When ranking using individual spectra, MVP demonstrates performance that is superior to or on par with existing methods, achieving 26.37% and 11.10% rank@1 for candidates by mass and formula, respectively. MVP offers a flexible, extensible foundation for learning from multiple molecule/spectra data views.

**For Table of Contents Only:** 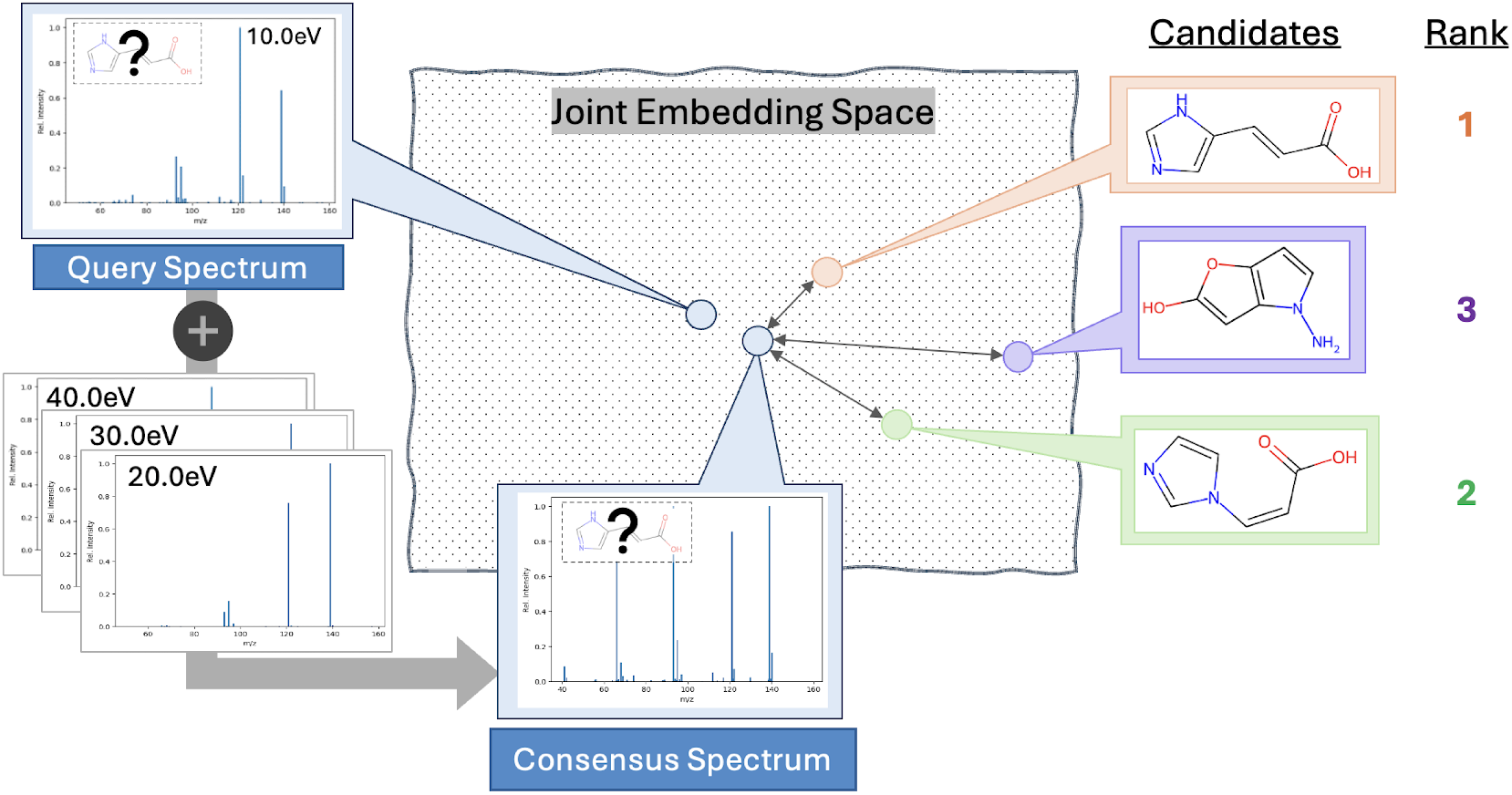

## Introduction

Untargeted metabolomics enables high-throughput profiling of thousands of metabolites, providing insights into cellular metabolism, disease mechanisms, biomarkers, and environmental effects. A fundamental challenge is metabolite annotation, the assignment of chemical structures to LC-MS/MS spectra. These spectra capture patterns of ion fragments as mass-to-charge (m/z) ratios and their corresponding intensities. Annotation typically involves matching experimental spectra to reference libraries such as NIST,^1^ GNPS,^2^ or MoNA.^3^ However, annotation rates remain low due to limited library coverage.^4^Low annotation rates have promoted the development of machine-learning (ML) methods that rank candidate molecules by their likelihood of producing a query spectrum. Existing approaches either map between molecular and spectral modalities or learn a joint embedding space. Mapping approaches include “inverse” methods such as de novo generation^5–8^ and fingerprint prediction^9,10^ (Figure 1A), and “forward” methods that simulate spectra from molecules^11–16^ (Figure 1B). In contrast, alignment approaches^17,18^ learn a joint embedding between the two modalities where matching molecule–spectrum pairs are placed close together, and non-matching pairs are positioned farther apart (Figure 1C). Alignment approaches, when implemented well, have shown to outperform mapping approaches, e.g., JESTR.^17^

**Figure 1.**
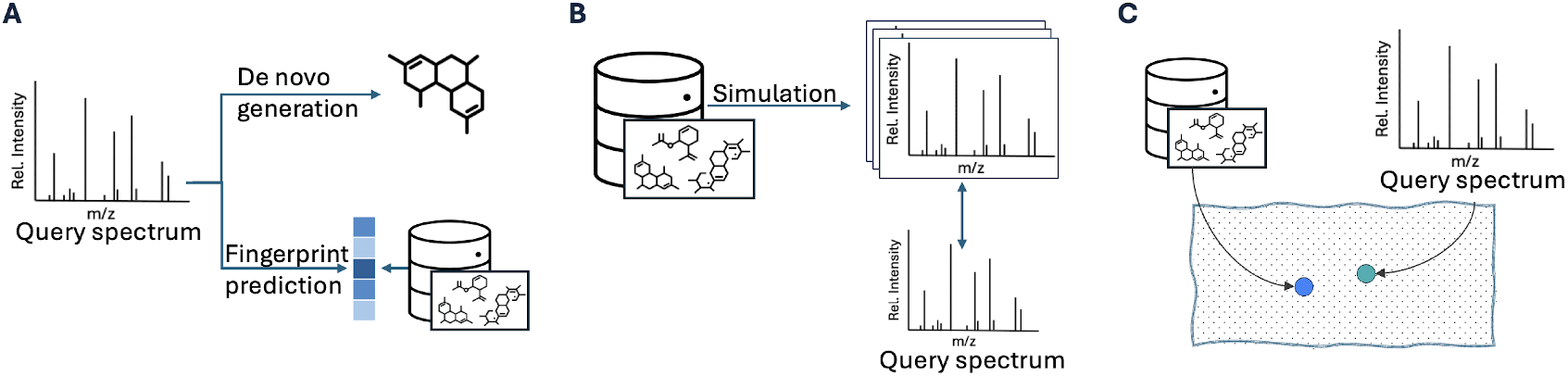
Three paradigms for spectra annotation: A) Inverse methods include de novo generation and fingerprint prediction from the query spectrum, B) Forward methods predict spectra from molecular candidates to match against the query spectrum, C) View alignment methods minimize the distance between a molecule and its spectral embedding.

Current alignment models, however, explore only a subset of the available views for each modality (Figure 2). Molecules can be represented as SMILES^19^/SELFIES,^20^ fingerprints,^21^ learned embeddings from pretrained models,^22,23^ structure-based fragmentation trees,^24^ chemical formulas, or 3-D structures. Spectra can likewise be described by spectral motifs,^25^ formula-based fragmentation trees, ^9^ subformula labels,^10^ learned embeddings from pretrained models,^26–28^ substructure labels, or consensus spectra. We hypothesize that combining multiple views enables the model to capture both complementary and shared information while mitigating view-specific artifacts, ultimately improving cross-modal alignment.

**Figure 2.**
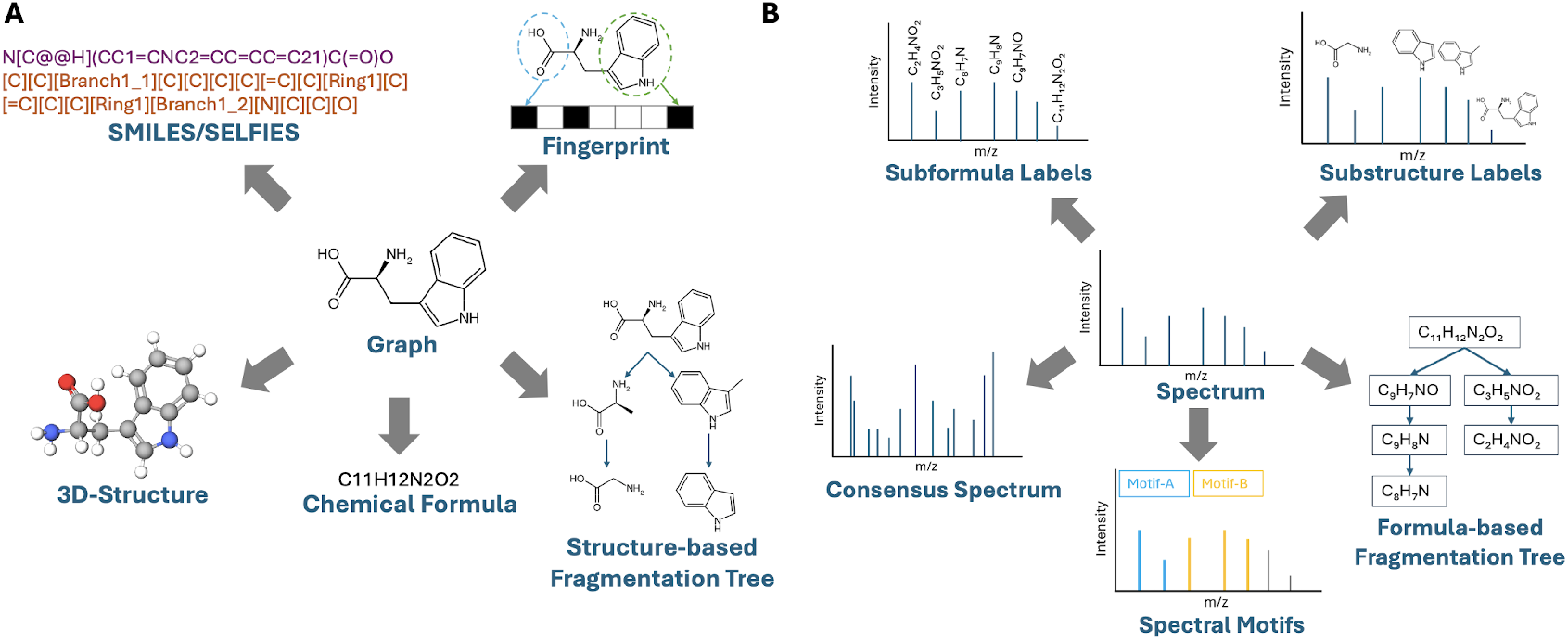
Different views of molecular data (A) and spectral data (B). The views at the center represent the most widely used views for metabolite annotation: spectrum and graph. Other representations not shown include latent embeddings from pretrained models for both molecules and spectra.

We present MultiView Projection (MVP), a contrastive framework that learns from multiple molecular and spectral views. Molecules are represented as graphs and fingerprints, while spectra are represented as both individual and consensus spectra (aggregated from multiple measurements). Consensus spectra, in particular, offer richer representations by aggregating measurements across collision energies, a common practice for improving spectrum quality.^29,30^ Prior studies have shown that using multiple spectra enhances annotation accuracy.^6,12,31^ Importantly, by maximizing mutual information across these four “training views”, MVP jointly aligns all views in a shared embedding space, enabling more robust candidate ranking. While some prior methods incorporate additional molecular or spectral views through concatenation or auxiliary prediction tasks (Section S1), MVP differs as it offers a systematic framework that unifies multiple views.

MVP employs four encoders, one per view, and optimizes a contrastive loss across all pairwise view combinations (Figure 3). At inference, it is possible to annotate data through one of many possible “ranking views”: graph-spectra (mol-s), fingerprint-spectra (fp-s), graph-consensus spectra (mol-cs), and fingerprint-consensus spectra (fp-cs). We evaluate two strategies for leveraging multiple spectra per molecule: an aggregate-then-rank strategy, which combines spectral information prior to candidate ranking, and a rank-then-aggregate strategy, which first ranks candidates for each individual spectrum and then aggregates the rankings. We train and evaluate MVP on the MassSpecGym benchmark,^32^ comparing against forward, inverse, and alignment-based baselines, ESP,^15^ MIST,^10^ and JESTR,^17^ respectively. Results show that MVP significantly improves candidate retrieval, particularly when leveraging consensus spectra. Ablation studies further validate the benefit of integrating all four modalities and employing the aggregate-then-rank strategy.

**Figure 3.**
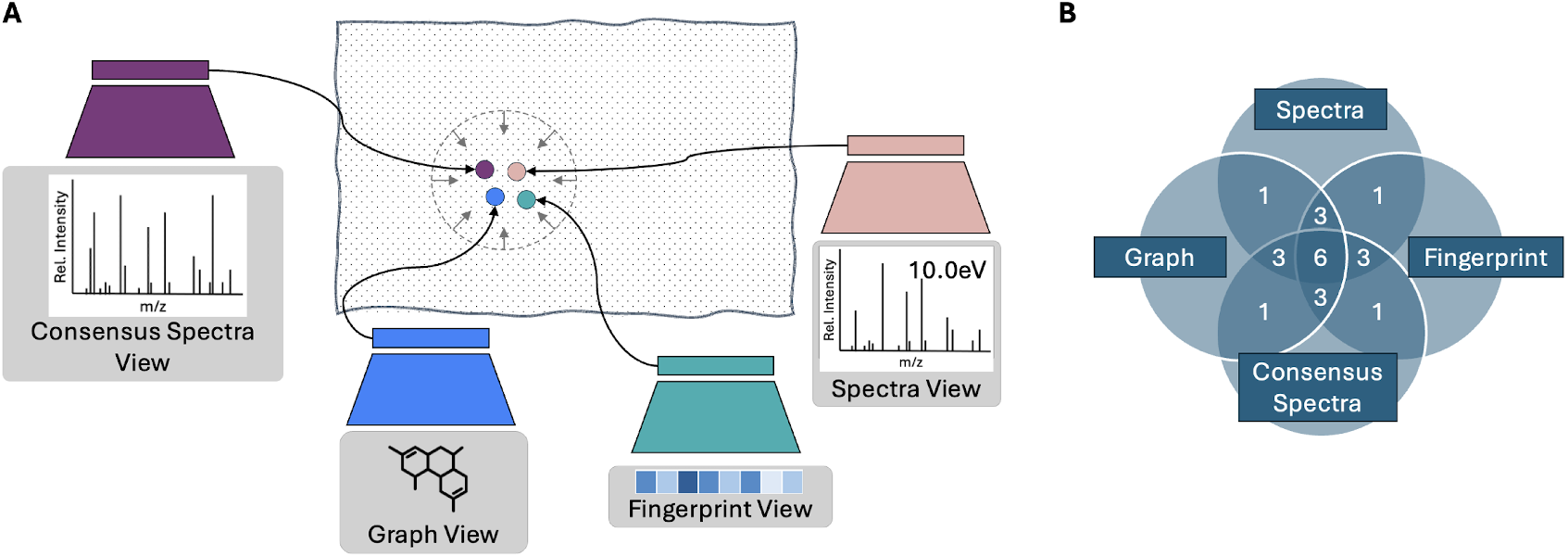
Overview of MVP. A) Four encoders are trained to place the four matching views close in the joint embedding space. B) MVP employs the full-graph paradigm, where the overlap of any number of views denotes the number of view pairs considered during training. For example, considering all four views results in six view pairs used for training.

Our contributions are as follows:

- Presenting MVP, a novel framework for jointly learning the embedding space of molecules and spectra that leverages four data views: molecular graph, molecular fingerprint, spectrum, and consensus spectrum. MVP learns representations that maximize the mutual information between different views of the same datum. We utilize and evaluate the learned representations for ranking molecular candidates for a given query spectrum.
- Achieving competitive results on the MassSpecGym retrieval task when candidates are provided by mass or by formula. In particular, for candidates by mass, MVP outperforms MIST by 145.83% and JESTR by 130.71% for rank@1. Further, our ablation studies demonstrate the value of jointly learning from additional spectra and molecule views beyond traditional spectrum/graph views used in prior work.
- MVP presents a systematic framework for effectively leveraging consensus spectra in molecular annotation. Our results consistently show that utilizing a consensus spectrum yields superior annotation results when compared to using a single spectrum. Further, we show that molecular annotation based on the consensus spectrum outperforms the aggregation of annotation results on individual spectra of the same molecule. That is, an aggregate-then-rank annotation paradigm is superior to a rank-then-aggregate paradigm.

## Methods

### Molecule and spectra view representations

MVP uses two molecule-based views, graphs and fingerprints. A graph encodes a molecular structure by treating atoms as nodes and bonds as edges. We use the same graph featurizer as JESTR,^17^ where node features include atom type, atomic mass, valence, ring membership, formal charge, radical electrons, chirality, degree, number of hydrogens, and aromaticity, and edge features include the bond type, ring membership, conjugacity, and the stereo configuration. A molecule’s fingerprint is attained using binary Morgan fingerprint (nbit=1024, r=5), computed using RDKit.^33^

MVP uses two spectra-based views, individual spectra and the consensus spectra. As with MIST, spectral peaks are first annotated using MIST’s peak formula annotation code based on an enumeration strategy given the ground truth chemical formula, keeping the most abundant 60 peaks for a given spectrum while using a *±* 20 ppm threshold between a peak’sm/z and a formula. Further, each peak is encoded as a vector *p ∈ ℛ*^15^, where the first 14 entries represent the atom count of elements, *E* = {C, H,O, N, P, S, Cl, F, Br, I, B, As, Si,Se} and the last entry represents the normalized intensity. The atom counts are scaled by dividing by the highest observed count of the corresponding element in the dataset. As the normalization depends on the characteristics of the training data, the model performance may not be optimal when a formula contains an atom that has not been seen or has a count value greater than that seen in the training data. Both spectra and consensus spectra are represented as a set of peaks, where each peak is defined using its formula and intensity.

For training, validation, and test sets, the consensus spectra are created by merging all peaks from spectra that share the same molecular identity. If multiple peaks share the same formula, one formula is kept, and the maximum of the intensities is retained.

### Encoders

MVP uses the same graph encoder as JESTR - a graph convolution network (GCN) of 3 layers followed by a pooling layer and an MLP with 2 fully connected layers to generate the final structural embeddings, *Z*_*mol*_ (Figure S1). The fingerprints are encoded by a 3-layered MLP to generate the fingerprint embeddings, *Z*_*fp*_. Both spectra and consensus spectra use encoders of the same architecture (Figure S2). Peaks are first encoded by a 3-layered MLP, and then the set of peaks is fed into a transformer encoder consisting of 2-attention heads, outputting embeddings of spectra *Z*_*s*_ and consensus spectra *Z*_*cs*_. The exact dimensions of the model architecture are described in Table S1. The model is trained with the Adam optimizer^34^ for 1,500 epochs with a learning rate of 7.0e-05, a batch size of 64, and a contrastive loss temperature of 0.05.

### Contrastive learning with multiple views

MVP contrasts four views of a datum: its molecular graph and fingerprint, spectrum, and consensus spectrum. The objective of contrastive learning is to place views of the same data point close in a shared embedding space while ensuring non-matching views are far apart. A non-matching set of views arises from a molecule and any non-matching spectra. We consider pairwise mutual information between the four views: *Z*_*mol*_, *Z*_*fp*_, *Z*_*s*_, and *Z*_*cs*_. The total loss function is formulated as:

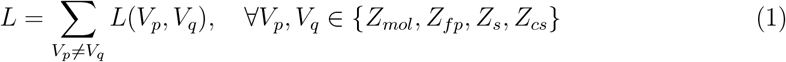

Each pairwise contrastive loss is computed symmetrically:

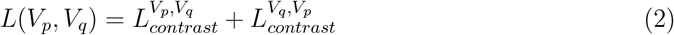

Following the CMC framework, the contrastive loss for a given view pair is defined as:

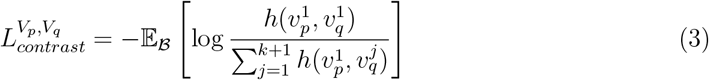

where 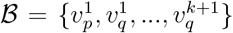 is a batch of samples, *k* is the number of negative samples 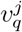 for a given sample 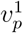, and 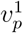 and 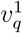correspond to a matching pair. The function *h* is a discriminatory function that measures the similarity between two embeddings using cosine similarity, scaled by a hyperparameter, *τ*. The hyperparameter adjusts how sharply or softly the model distinguishes between positive and negative samples, with smaller *τ* encouraging the model to push negative Pairs further apart. The function *h* is defined as:

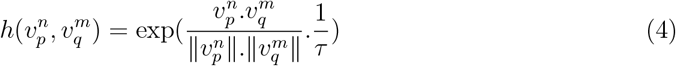

## Results

MVP is evaluated on the retrieval task, where given a spectrum and a set of molecular candidates, MVP ranks the molecules using the embedding similarity of a spectra-based view against a molecule-based view. Consistent with standard benchmarking metrics, we report rank@k, where *k* = 1, 5, 20 indicates the percentage of the test data for which the correct molecule is ranked in the top *k* position. When we compare MVP against other annotation tools, we also report the MCES@1, a metric that measures the maximum common edge subgraph (MCES) distance between the target molecule and the molecule ranked at 1. The best-performing model achieves the highest rank@k scores and the lowest MCES@1 value.

### Dataset

We evaluate MVP on the MassSpecGym dataset, a recently established benchmark dataset for machine learning models for annotation-related tasks. Two candidate sets are provided. Candidates by mass are molecules whose molecular mass is within 10 ppm of that of the target molecule, while candidates by formula are molecules with the exact chemical composition as the target. Each molecule is associated with a maximum of 256 molecular candidates for each candidate set. MassSpecGym contains spectra with [M+H]+ and [M+Na]+ adducts. Peak labels were obtained through MIST’s formula annotation algorithm, which assumes that every peak is charged with the adduct, an assumption that may not always hold. Under this framework, peaks are labeled without the adduct but can be merged across spectra with different adducts if they share the same molecular formula. This facilitates the construction of the consensus spectra, though it does not account for possible differences in ion response between adducts. To mitigate potential inaccuracies from these assumptions, we assess performance not only on the whole dataset (Table 1A) but also on the [M+H]+ subset (Table 1B) in some of our evaluations. By design, the MassSpecGym dataset is challenging as the spectra in the dataset are split such that molecules across splits have an MCES distance of at least 10 from one another. Strong performance on the MassSpecGym indicates that models exhibit strong generalization and robustness. MassSpecGym benchmark provides a reproducible and fair comparison with existing state-of-the-art methods, which were previously trained on this benchmark set.

**Table 1:**
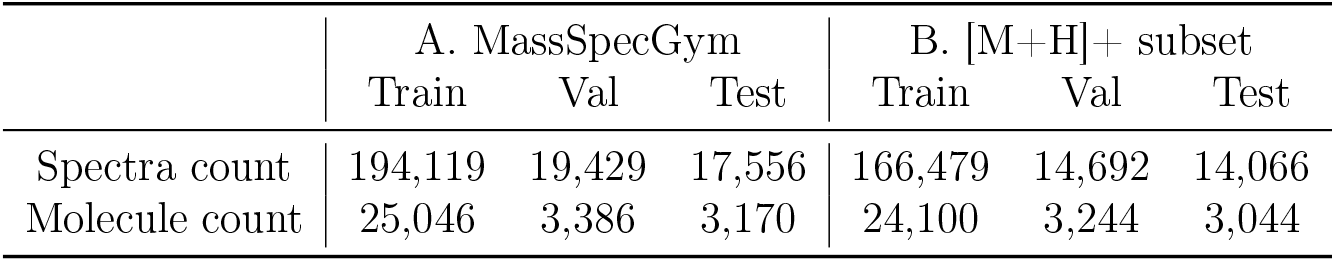
MassSpecGym statistics on the whole dataset (A) and the [M+H]+ subset (B).

### Which ranking view is best?

Across all ranking views, the best performance is consistently attained using the mol-cs ranking view for both candidates by mass and candidates by formula (Table S2). The performance improvements across the ranks when using the consensus spectra (mol-cs vs mol-s, and fp-cs vs fp-s) characterize the significant impact of leveraging multiple fragmentation patterns within each consensus spectrum at inference. The results indicate that MVP is able to place the consensus spectrum, on average, closer to the molecular views compared to individual spectra. Using the graph view provides better ranking performance than the fingerprint view. That is, mol-s ranking view always outperforms fp-s, and mol-cs outperforms fp-cs. The graph view, therefore, likely shares more mutual information with a spectrum. For the rest of this paper, we focus further evaluation on ranking views that use the graph view, but highlight the importance of the fingerprint view in the ablation study on the various training views.

### Comparison with other annotation tools

We compare MVP against MIST, ESP, and JESTR (Table 2). We also provide partial results for the MassSpecGym dataset on SIRIUS^35^ (Section S3.2), as its implementation is not publicly available and it cannot be pretrained on the same dataset. MVP with the mol-cs ranking view outperforms other tools across all metrics for both candidates by mass and formula. Compared to MIST, MVP with the mol-cs ranking view improves rank@1 by 145.83% (candidates by mass) and 45.87% (candidates by formula), and MCES@1 by 39.36% (mass) and 18.67% (formula). Compared to JESTR and JESTR_NR_, MVP achieves average gains of 134.29% (mass) and 17.96% (formula) in rank@1, and 54.18% (mass) and 11.33% (formula) in MCES@1. Using the mol-s ranking view, MVP outperforms other works except for rank@1 and rank@5 compared to JESTR when ranking candidates by formula. It is possible that MVP’s use of peak formulae as input to the spectral encoders results in poor differentiation of molecular candidates, possibly explaining MVP’s worse performance than JESTR, which does not utilize peak formulae. We further explore this issue in the comparative study on spectral representation. Nevertheless, the MCES@1 for MVP (mol-s) is smaller than that of JESTR, suggesting that, in general, candidates ranked in the first position by MVP are more similar to the target structures than those ranked by JESTR. Overall, the improvement is more significant in ranking candidates by mass than in ranking candidates by formula, likely because the formula-annotated peaks help discern candidates of the same mass but different formulae.

**Table 2:** Ranking Performance on MassSpecGym compared to other tools. The best performance is **highlighted**, and the second best performance is underlined. The values in brackets indicate 99.99% confidence intervals upon bootstrapping (20,000 resamples).

| Model | rank@1 (↑) | rank@5 (↑) | rank@20 (↑) | MCES@1(↓) |
| --- | --- | --- | --- | --- |
| A. Candidates by Mass |  |  |  |  |
| MIST | 14.64 (13.82-15.54) | 34.87 (33.69-36.10) | 59.15 (57.89-60.39) | 15.37 (15.12-15.32) |
| ESP | 10.71 (9.95-11.52) | 24.84 (23.76-25.92) | 42.66 (41.51-43.89) | 35.36 (34.94-35.81) |
| JESTR | 15.13 (14.27-16.04) | 36.75 (35.61-37.95) | 60.32 (59.13-61.49) | 20.71 (20.32-21.13) |
| JESTR<sub>NR</sub> | 15.60 (14.71-16.48) | 37.47 (36.28-38.64) | 60.55 (59.39-61.74) | 19.98 (19.61-20.35) |
| MVP (mol-s) | 26.37 (25.25-27.46) | 58.90 (57.73-60.13) | 86.88 (86.05-87.69) | 12.27 (11.98-12.58) |
| MVP (mol-cs) | **35.99 (34.82-37.20)** | **67.97 (66.81-69.06)** | **91.92 (91.22-92.58)** | **9.32 (9.09-9.56)** |
| B. Candidates by Formula |  |  |  |  |
| MIST | 9.57 (8.88-10.30) | 22.11 (21.10-23.13) | 41.12 (39.98-42.34) | 12.75 (12.59-12.91) |
| ESP | 11.05 (10.28-11.82) | 27.42 (26.36-28.53) | 52.20 (51.00-53.46) | 16.44 (16.30-16.59) |
| JESTR | 11.85 (11.06-12.66) | 32.95 (31.80-34.14) | 61.46 (60.23-62.71) | 11.68 (11.51-11.86) |
| JESTR<sub>NR</sub> | 11.82 (11.05-12.68) | 33.48 (32.31-34.68) | 61.46 (60.22-62.72) | 11.71 (11.53-11.88) |
| MVP (mol-s) | 11.10 (10.37-11.88) | 31.14 (30.00-32.28) | 61.99 (60.82-63.20) | 11.61 (11.44-11.77) |
| MVP (mol-cs) | **13.96 (13.15-14.83)** | **36.88 (35.70-38.04)** | **68.12 (67.00-69.23)** | **10.37 (10.20-10.53)** |

### Aggregate-then-rank using the consensus spectrum or rank-then-aggregate?

Given multiple query spectra measurements for the same molecule and a set of candidate molecules, MVP provides the opportunity to first create a consensus spectrum and then rank the candidates using the mol-cs ranking view. MVP therefore implements an “aggregate-then-rank” paradigm. As an alternative, it is possible to rank the candidates for each spectrum and then aggregate the ranking results. Within this “rank-then-aggregate paradigm”,

MVP provides a ranking for each candidate molecule for each query spectrum using the mol-s ranking view. As such a strategy suggests different rankings on the candidates for each query spectrum, we evaluate two rank aggregation methods. The *average rank* method averages the rank of each molecular candidate across all spectral queries. The ranking of each candidate is based on its mean rank, with lower average ranks indicating a better rank. This simple and intuitive method equally weights all spectral queries. The *reciprocal rank* method assigns a score by summing the reciprocal of the ranking position (1/rank) for each candidate. A candidate with a higher score ranks better. This method prioritizes highly ranked candidates more strongly, making it more sensitive to cases where a candidate is ranked very well in some queries but poorly in others. An illustration of both paradigms is shown in Figure S3.

We compare the two rank-then-aggregate methods against MVP’s mol-cs ranking. As the number of spectra per target molecule varies in the test set, we limit the evaluation to a subset of the test set for which a test molecule has exactly three spectra, since the majority of the molecules have three spectra. There are 1,531 such cases (Figure S4). The rank-then-aggregate methods underperform the aggregate-then-rank method (Figures S5A and S5B). MVP therefore provides an effective method for exploiting the additional coverage provided by spectra measured at different collision energies and/or detected with different adducts.

Like MVP, CFM-ID^12^ leverages multiple spectra for annotation. Users can input up to three spectra corresponding to low, medium, and high collision energies, which are then matched against simulated spectra. Candidate molecules are ranked based on the average score across the three simulated spectra, following a rank-then-aggregate approach. We benchmark against the publicly available CFM-ID 4.0 pretrained model using a merged spectra dataset derived from the MassSpecGym dataset, and ranking the candidates by formula. For each test molecule, we randomly sample three spectra (one for each collision energy level) and rank the candidates accordingly. To ensure a fair comparison, we exclude spectra whose molecules appear in CFM-ID’s training set. CFM-ID achieves a 2.00% rank@1 performance (60/3,004 consensus spectra), whereas MVP achieves 13.91% rank@1 (418/3,004), further demonstrating the advantages of the aggregate-then-rank strategy. When leveraging all available spectra for each molecule, MVP further improves to 14.38% rank@1 (432/3,004), emphasizing the value of incorporating all relevant information. The effectiveness of using the consensus spectrum depends on the availability of multiple spectra collected at various collision energies. Therefore, molecules with limited collision energy coverage may not benefit from using the consensus spectra view. MVP addresses this limitation with a flexible design that leverages available data to improve annotation accuracy.

### Comparative study on spectra representation

As JESTR, which uses binned spectral representation without formulae, outperformed MVP on candidates by formula for ranks 1-5, we compare peak formulae representation of the spectra with a binned representation. We denote the MVP spectra representation as FormSpec, whereas the binned version is referred to as BinnedSpec. The MVP architecture setup remains the same, except that the spectra and consensus spectra encoders are replaced with 3-layered MLPs, similar to the spectra encoder architecture used in JESTR (Figure S2). We compare MVP when trained on MassSpecGym and the [M+H]+ subset. Overall, using FormSpec provides improved results over the BinnedSpec across both datasets and all metrics, except for rank@1 for both mol-cs and mol-s ranking views and rank@6-8 for the mol-cs ranking view with formula-based candidates when trained on the whole dataset (Table S3 and staircase plots in Figures S6A and S6B).

For candidates by mass, when using both mol-s and mol-cs ranking views, there is a remarkable improvement across all ranks using the formula representation. The mol-s ranking view achieves 26.37% rank@1 with FormSpec vs 12.29% rank@1 with BinnedSpec, a 114.56% improvement. The mol-cs ranking view achieves 35.99% rank@1 with FormSpec vs 16.63% with BinnedSpec, a 116.42% improvement when evaluated on the whole dataset. These results indicate that MVP’s spectral encoders can clearly differentiate candidates from the target molecule. When analyzing the distribution of cosine similarities between the query spectrum and its target molecules against the average similarity of the query spectrum and the corresponding candidate molecules, MVP’s formula-based spectral encoders better differentiate the target molecule from the candidates when compared to using binned-based spectral encoders (Figures S7A vs B). The results when using the [M+H]+ subset are consistent with those obtained using all data points for the MassSpecGym.

For candidates by formula, training MVP using binned spectra outperforms the use of formula-based spectra for rank@1, but the FormSpec model outperforms the BinnedSpec at higher ranks (Figure S6B). When examining the distribution of the cosine similarities between query spectrum and target molecule and the average cosine similarity between query spectrum and candidate molecules, the differentiation between target and candidate is not as pronounced as was the case for candidates by mass (Figure S7C). The distribution of the candidate-spectra similarity shifts to the right, compared to the case when using FormSpec, along with the target-spectra similarity. This reflects the difficulty in distinguishing candidates from their relevant targets when they share the same chemical formulae. The right shift is more pronounced for FormSpec compared to BinnedSpec (Figure S7C vs D), indicating that the spectra and molecule encoders are learning associations related to the overall chemical composition of the molecule rather than the structural relation between spectra and molecules. However, when utilizing the [M+H]+ subset, FormSpec outperforms BinnedSpec, indicating that potentially inaccurate peak labels for the [M+Na]+ spectra may contribute to the poorer performance when using FormSpec at rank@1 when training on the MassSpecGym.

As there are multiple spectra per molecule in the test set, we examine the correlation of the two spectral representations with rank variability. For all test molecules with 2, 3, or 4 spectra, we compute the normalized rank difference, defined as the range of the ranks divided by the number of candidates (Figures S7E and S7F). For both candidates by mass and formula, we observe significantly smaller rank variability when using FormSpec, therefore indicating that the use of formulae enables the spectra encoder to learn a more uniform spectra representation for the same molecule.

### Are all four views needed? An ablation study on training views

We study the contribution of each view by eliminating one or more views when training the model, resulting in four models, trained on mol-fp-s-cs, mol-s-cs, mol-fp-s, or mol-s training views (Table S4A). There are no clear patterns of which of the four models consistently outperforms the others across all ranks. For example, when ranking candidates by mass using the mol-cs view, the mol-fp-s-cs and the mol-s-cs models are competitive across the ranks (Figure S8A). Similar competitive patterns are observed with the four models using the mol-s ranking view. When ranking candidates by formula, competitive patterns are also observed across the ranks (Figure S8B). These observations suggest that additional views do not consistently improve performance across the ranks.

However, when training only on the [M+H]+ subset of the MassSpecGym data (Table S4B), we observe more consistent patterns across the ranks. The mol-fp-s-cs model trained on all four views outperforms models trained on fewer views, demonstrating the value of additional views. Furthermore, removing the fingerprint view consistently reduces performance. For example, when ranking candidates by mass with the mol-s views, the rank@1 performance drops from 29.73% to 25.56% when the fingerprint view is removed from the mol-fp-s-cs model. Similarly, the rank@1 drops from 29.22% to 27.83% when the fingerprint view is removed from the mol-fp-s model. Similar drops in performance due to removing the fingerprint view are observed when ranking candidates by formula, indicating the fingerprint view offers nuances critical for ranking. Finally, considering that the performance of the models when trained on the [M+H]+ subset surpasses that of models trained on the whole dataset, training specialized models or finetuning a general model for each adduct may be beneficial.

### MVP reveals differential metabolites in cultured and conventional meat

To illustrate the applicability of MVP within metabolomics workflows, we apply it to data from a recent study comparing the metabolomic profiles of traditional chicken meat (CON), muscle satellite cells (CIC), and myotube-formed cells (CAC).^36^ This investigation is particularly important for evaluating the nutritional value and safety of cultured meat, an emerging alternative to conventional animal products. LC–MS/MS data in mzML format are publicly available on the Metabolomics Workbench.^37^ The original study identified 321 features corresponding to 214 unique molecules.

For this evaluation, we focus on the positive ion mode data. Since neither the processed spectra nor the mapping to the identified metabolites was provided, we replicated the study using a standard metabolomics workflow and report the hit rate. Feature extraction and representative spectra selection are performed using MZmine3,^38^ followed by formula and adduct assignment with SIRIUS v6.1.0.^35^ Candidate structures are retrieved within a 0.01 Da tolerance from FoodDB, ^39^ HMDB,^40^ MassSpecGym,^32^ NIST23,^1^ and the GNPS Multiplex library.^41^ Further, recognizing that MVP relies on the correct molecule being present in the candidate pool and accounting for potential adduct misassignments, we include all reported compounds in the candidate set. To improve confidence in annotations by MVP, we focus on results with a molecule–spectra score greater than 0.5 and a margin of at least 0.01 between the top- and second-ranked candidate scores.

Using this workflow, 900 features are detected with MZmine3, of which 866 receive formula and adduct assignments from SIRIUS. MVP subsequently annotates 640 spectra corresponding to 146 unique molecules that met the defined criteria (Figures S9A and S9B). Metabolite classification using NPClassifier^42^ demonstrates that MVP annotations encompass all pathways reported in the original study, with consistent enrichment patterns (Figures 4A and S9C). In particular, amino acids and peptides are significantly more abundant in conventional meat compared to cultured meat, while other metabolic pathways showed comparable abundances.

**Figure 4.**
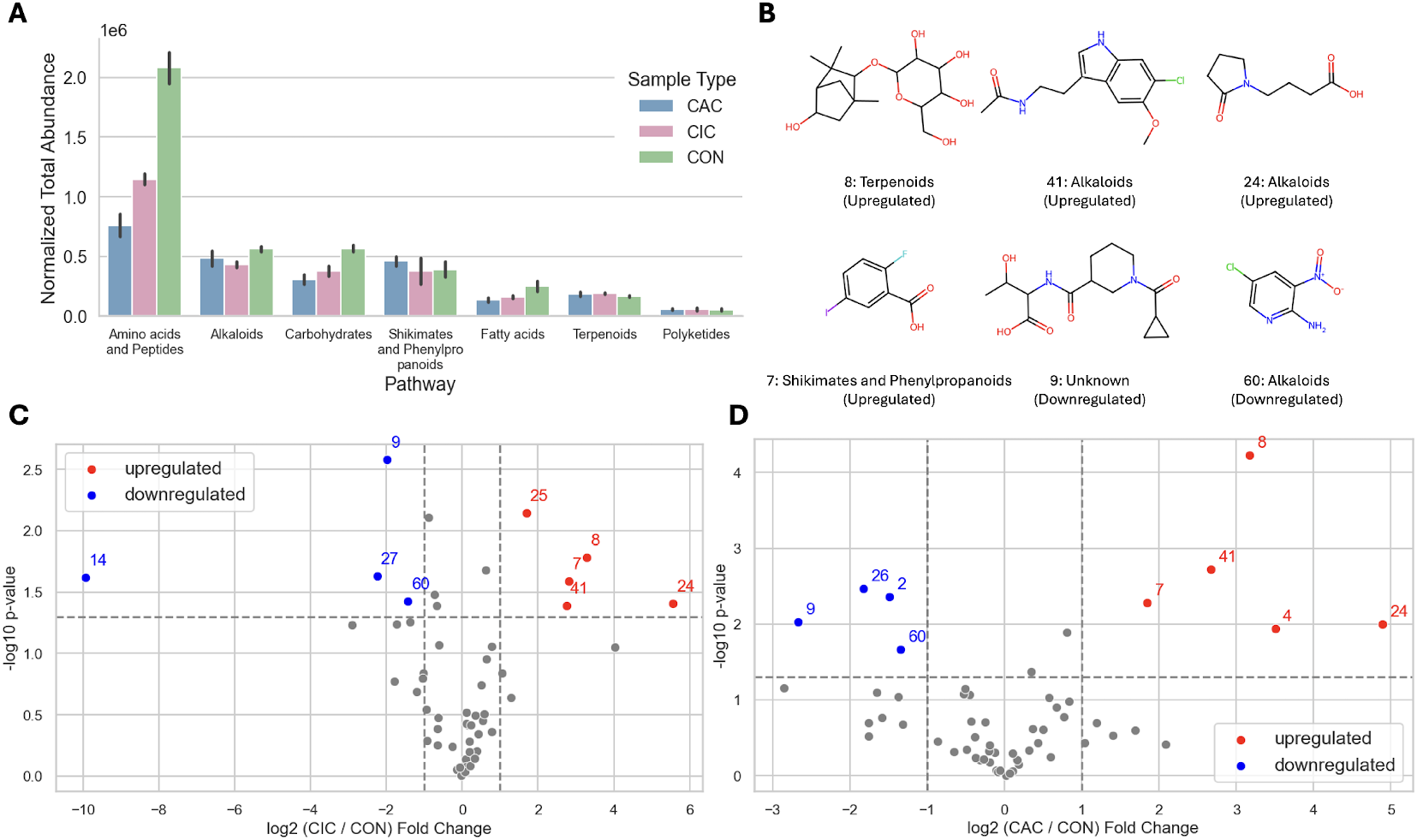
Case study results. A) Total abundance of each pathway per sample type. B) Putative metabolites identified by MVP. Each molecule is labeled with an ID, and its pathway and regulation. The IDs correspond to the labels in the volcano plots that show the fold change and p-value between CIC and CON features (C) and CAC and CON features (D).

Of the 146 metabolites identified by MVP, 73 are newly annotated, highlighting the model’s ability to expand metabolite coverage beyond existing workflows. Several metabolites (Figure 4B) that are associated with the terpenoid, alkaloid, and shikimate and phenylpropanoid pathways are always differently regulated in both CIC and CAC relative to CON (Figure 4C and 4D). Four molecules (#7, 8, 24, and 41) are consistently upregulated, and two (#9 and 60) are downregulated. The reported annotations are putative and require experimental validation.

## Conclusion

We presented MVP, a novel framework for learning a joint spectral/molecular embedding space using multiple data modalities: molecular graphs, fingerprints, spectra, and consensus spectra We demonstrated the effectiveness of this framework in ranking molecular candidates for spectral annotation. While prior work incorporated these modalities as multi-tasks or via concatenation, MVP jointly learns from the mutual information among the views. Another contribution of this work is that MVP is the first annotation tool designed to enable annotation based on the consensus spectra. While each individual constituent spectrum is a subset of the consensus spectrum, the consensus spectrum view serves as an anchor for different spectra of the same molecule, promoting the model to learn uniform spectra representation regardless of instrument conditions when they share the same molecule. Comparing annotation using the consensus spectrum against ranking-then-aggregating ranking results for its constituent spectra, we showed that the aggregate-then-rank paradigm produces superior performance. We also showed that annotation using the mol-cs view outperforms all other views. On the MassSpecGym benchmarking dataset, MVP’s mol-cs ranking view achieves 35.99% rank@1, a 145.83% and 134.29% improvement over MIST and JESTR, respectively, for candidates by mass. For candidates by formula, MVP achieves 13.96% rank@1, a 45.87% and 17.96% improvement over MIST and JESTR, respectively. The ability to rank candidate molecules based on the query consensus spectrum will be valuable for practitioners who often extract several measurements for the same metabolite. With MVP, there is no need to decide how to aggregate and interpret the rankings for the various individual constituent spectra. Further, our comparative and ablation studies show that including peak formulae is advantageous for ranking candidates by mass, and that learning from all four views provides improved performance compared to fewer views. Lastly, we demonstrate a practical integration of MVP with existing metabolomics workflow on a study that compares metabolic profiles between cultured and conventional meat. With MVP annotations, we observe underrepresented amino acids and peptides in cultured meat, consistent with the original findings. Additionally, 73 putative metabolites are uncovered, with six consistently up or downregulated in cultured meat samples. Overall, MVP demonstrates the potential of using additional information beyond a spectrum and its corresponding molecular structure to improve representation quality for molecule-spectra matching. While MVP achieves strong performance, there are several directions for future improvement. Representation quality could potentially benefit from more accurate peak labeling and additional views. As ranking is a separate task from learning representations, including losses that prioritize ranking molecular candidates could improve ranking performance. Finally, MVP can be extended to related tasks such as ranking scaffolds and identifying analogs, further demonstrating its versatility and impact.

## Data availability

The data, model scripts, and pretrained models underlying this study are available at https://github.com/HassounLab/MVP.

Web Interface: We developed a web-based interface for MVP, hosted on Hugging Face Spaces, to allow researchers to easily explore the tool. Users can upload an MGF file and a candidate JSON file to perform annotation. We provide sample files for convenience. Upon completion, the interface provides annotation results in a downloadable CSV file. As the web application runs on CPU resources, we limit the number of molecule-spectra pairs that can be run per instance. For large-scale datasets, we recommend users clone our repository and run MVP locally on GPU-enabled machines. The tool is available at https://huggingface.co/spaces/HassounLab/MVP.

## Supporting Information

Comparison to other tools, model architecture design, ablation study results, comparison against SIRIUS, aggregate-then-rank vs rank-then-aggregate experiment setup and results, comparative study on spectra features, and biological case study.

## Funding

Research reported in this publication was supported by the National Institute of General Medical Sciences of the National Institutes of Health under award number R35GM148219. The content is solely the responsibility of the authors and does not necessarily represent the official views of the NIH.

## Notes

### Competing Interest Statement

The authors have declared no competing interest.

