## Supporting Information for "Learning from All Views: A Multiview Contrastive Framework for Metabolite Annotation"

#### S1 Comparison to other tools

MVP differs from existing methods that incorporate additional molecular or spectral modalities. Below, we highlight key distinctions.

**Molecular representations**: ESP[S1] trains separate graph and fingerprint encoders for spectrum prediction and merges their outputs, with spectral motif prediction as an auxiliary task. CMSSP[S2] concatenates graph and fingerprint representations into a joint embedding. MIST[S3] uses spectra labeled with sub-formulae and predicts substructure-level fingerprints. These approaches show that enriched modalities (e.g., labeled spectra) can improve performance and that auxiliary tasks aid generalization. However, they often rely on ad hoc concatenation or multitask strategies rather than a systematic framework. In contrast, MVP jointly aligns all views in a shared embedding space, capturing complementary and shared information without requiring auxiliary prediction tasks.

**Spectral representations**: Several methods incorporate multiple spectra per molecule. CFM-ID[S4] simulates spectra at different collision energies and compares them to multiple experimental spectra. Spec2Mol[S5] uses a CNN with four input channels to learn from spectra measured under different precursor ions and energies. Such designs are effective when datasets are uniformly structured but often fail in practice, where the number and type of spectra vary. MVP addresses this limitation by aggregating available spectra into a consensus spectrum, ensuring that no information is discarded or underutilized.

**Consensus spectra**: ChemEmbed[S6] trains a consensus spectrum encoder to predict Mol2Vec embeddings,[S7] demonstrating that consensus spectra improve learned embeddings. However, this approach is limited to consensus inputs and cannot handle single-spectrum queries. MVP introduces dedicated encoders for both individual and consensus spectra, making it more flexible and applicable across diverse datasets.

**Methodological extensions**: Conceptually, MVP extends JESTR,[S8] which learns a joint embedding of molecular graphs and spectra using contrastive multiview coding (CMC). MVP expands this to four views (graph, fingerprint, spectrum, consensus spectrum), updates the spectral encoder from an MLP to a transformer (as in MIST), and represents spectra as sets of formulae rather than fixed-dimension vectors.

In summary, prior approaches leverage additional modalities or multiple spectra in ad hoc ways, whereas MVP provides a systematic framework for learning from multiple molecular and spectral views, with explicit support for both individual and consensus spectra.

### S2 Method Details

#### S2.1 Molecule encoder

The molecule encoder is depicted in Figure S1 and further described in Table S1A.

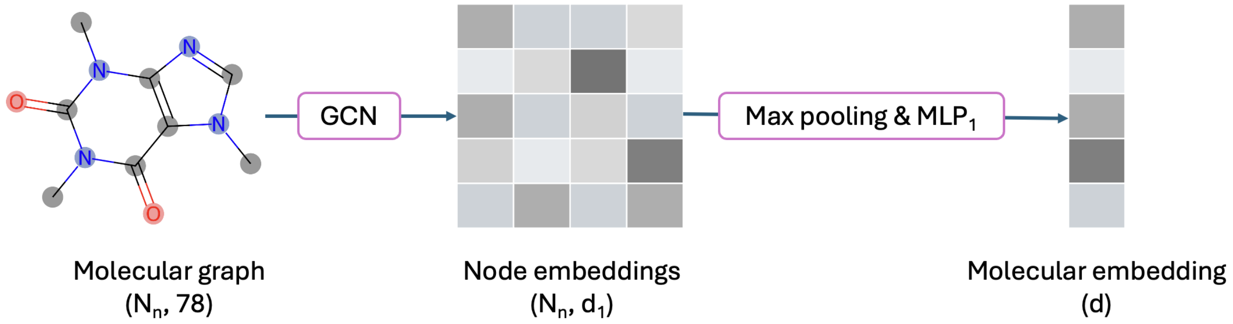


Figure S1: Architecture of molecule encoder. A molecule is encoded as a graph with $N_n$ nodes and embedded using a graph convolution network (GCN). Node embeddings are aggregated using max pooling before passing through $MLP_1$ to obtain the final molecular embedding. Parentheses indicate the dimensionality of tensors at each stage.

#### S2.2 Spectra encoders

MVP represents spectra as a FormSpec (Figure S2 top and Table S1B), in contrast to a BinnedSpec used by JESTR[S8] (Figure S2 bottom and Table S1C).

### S3 Additional Results

#### S3.1 Performance with various ranking views

MVP is trained on four views (two molecule-based views: mol and fp, two spectra-based views: s and cs). During inference, the cosine similarity between a spectra-based view and a molecule-based view is used for ranking. There are four such combinations: mol-s, fp-s, mol-cs, fp-cs (Table S2).

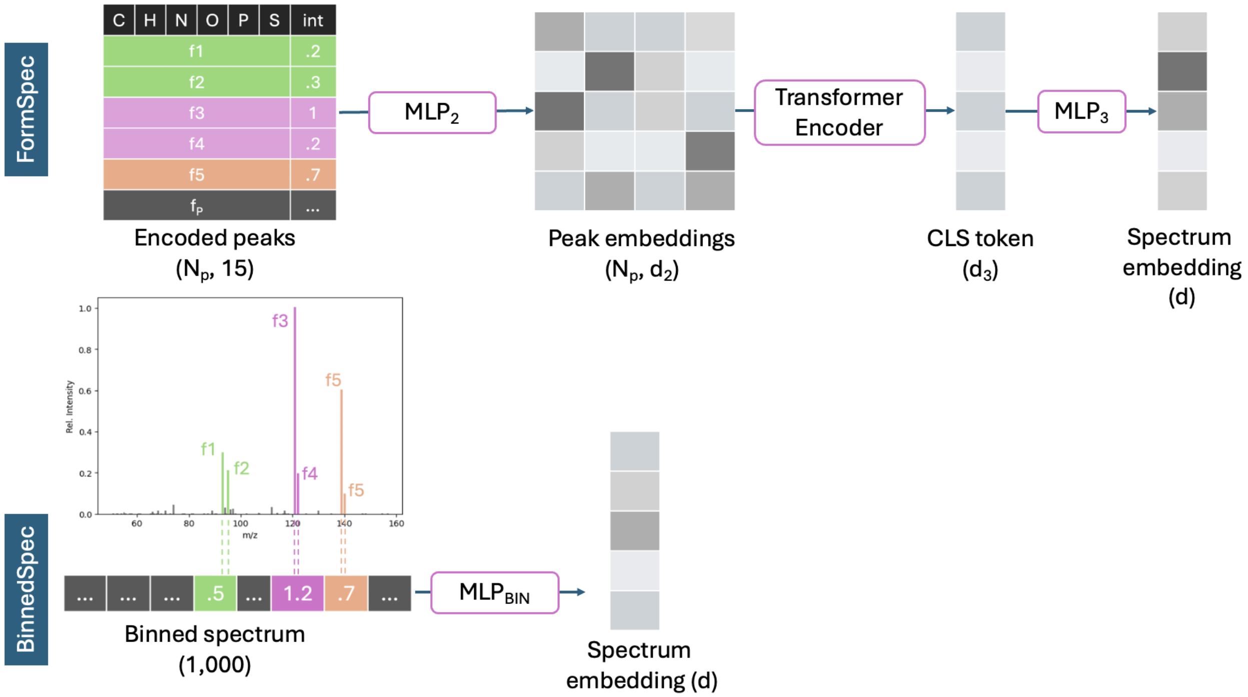


Figure S2: Architecture of spectra encoders. Top: FormSpec encodes $N_p$ peaks as a set of molecular formulae ($f_1 ... f_p$), concatenated with their peak intensities (int). Peaks are embedded using $MLP_2$, then fed into the transformer encoder. The [CLS] token of the spectrum is passed through $MLP_3$ to obtain the final spectrum embedding. Bottom: BinnedSpec bins the spectrum into a vector of 1,000 dimensions before passing through $MLP_{BIN}$ to obtain the final spectrum embedding. Parentheses indicate the dimensionality of tensors at each stage.

Table S1: Hyperparameters for A) molecule encoder, B) FormSpec encoder, C) BinnedSpec encoder

| A. Molecule encoder | |
|---|---|
| GCN | Input dimension: 78<br>Hidden dimensions: [64, 128, 256]<br>Dropout: 0.3<br>Pooling: Max pooling |
| $MLP_1$ | Input dimension: 256<br>Hidden dimensions: [1024, 512]<br>Activation function: ReLU |
| B. FormSpec encoder | |
| $MLP_2$ | Input dimension: 16<br>Hidden dimensions: $[64, 128, 256]$<br>Dropout: 0.2<br>Activation function: ReLU |
| Transformer encoder | Dim: 256<br>Num layers: 2<br>Aggregation: [CLS] token |
| $MLP_3$ | Input dimension: 256<br>Hidden dimensions: [512]<br>Activation function: ReLU |
| C. BinnedSpec encoder | |
| $MLP_{BIN}$ | Input dimension: 1000<br>Hidden dimensions: [512, 1024, 1024, 512]<br>Activation function: ReLU |

Table S2: MVP performance using different ranking views. The best performance is **highlighted**, and the best performance based on the spectra view is underlined.

| Ranking views | rank@1 (↑) | rank@5 (↑) | rank@20 (↑) |
|---|---|---|---|
| A. Candidates by Mass | | | |
| mol-s | 26.37 | 58.897 | 86.88 |
| fp-s | 15.48 | 36.02 | 62.73 |
| mol-cs | **35.99** | **67.97** | **91.92** |
| fp-cs | 23.27 | 46.09 | 71.66 |
| B. Candidates by Formula | | | |
| mol-s | 11.10 | 31.14 | 61.99 |
| fp-s | 9.08 | 22.00 | 43.11 |
| mol-cs | **13.96** | **36.88** | **68.12** |
| fp-cs | 12.99 | 25.31 | 47.94 |

#### S3.2 Comparison against SIRIUS

We benchmark MVP against the publicly available SIRIUS software on the MassSpecGym dataset, using candidates by formula. Since SIRIUS is not trained on the same dataset used by MVP, we report the performance on the subset of test spectra (12,314/17,556) where the target molecules are not in SIRIUS's training set (`https://csi.bright-giant.com/v3.0/api/fingerid/trainingstructures?predictor=1`). Using the SIRIUS graphical user interface (version 6.1.0), we created a custom database containing all candidates in the test dataset. SIRIUS correctly annotates 1,281 spectra, achieving a 10.41% rank@1, while MVP mol-cs achieves 16.70%, a 60.42% relative gain, and MVP mol-s achieves 12.72%, a 22.19% relative gain. On the complete test set, SIRIUS correctly ranks the target molecule in 3,342 cases (19.04% rank@1 vs 13.96% for MVP mol-cs and 11.10% for MVP mol-s). These results show that MVP generalizes beyond its training data more effectively than SIRIUS.

#### S3.3 Aggregate-then-rank vs Rank-then-aggregate

A molecule is often measured at various collision energies with various adducts. To utilize a trained model, practitioners can either rank a set of candidates for each individual spectrum and aggregate the ranking results (rank-then-aggregate) or merge the spectra into a single consensus spectrum and then rank a set of candidates (aggregate-then-rank) (Figure S3). This experiment utilizes the subset of test molecules with exactly three spectra in the test set (Figure S4). Aggregate-then-rank using the mol-cs view outperforms rank-then-aggregate approaches that use the mol-s view (Figure S5).

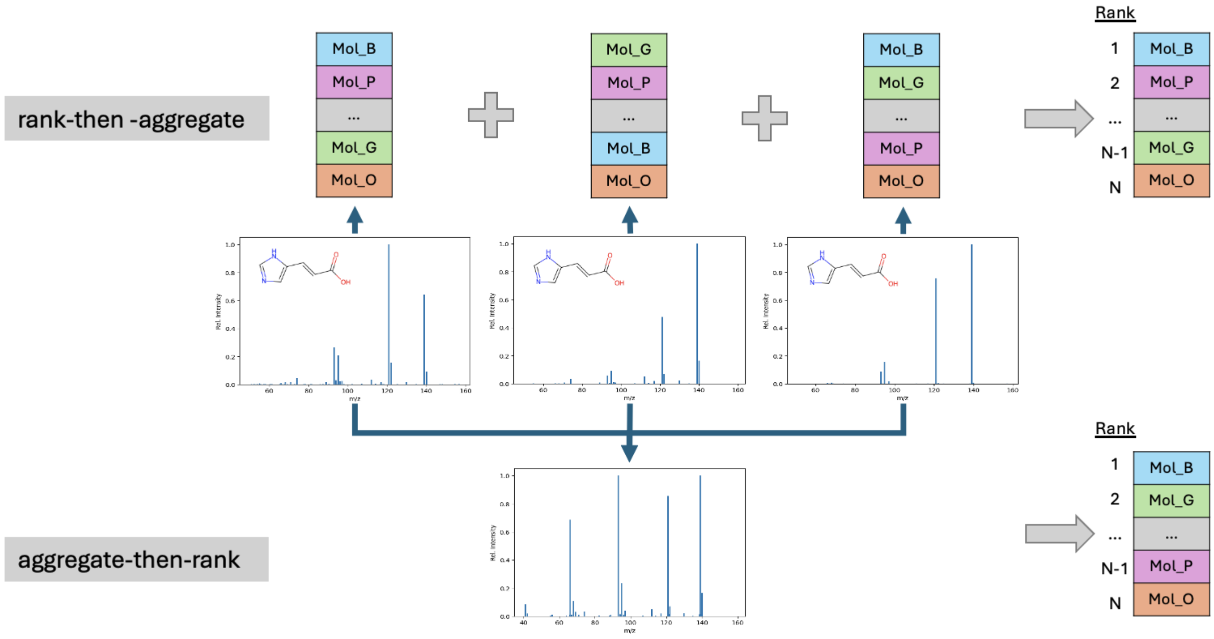


Figure S3: Overview of the rank-then-aggregate and aggregate-then-rank methods. For rank-then-aggregate methods, candidates for each spectrum are ranked, then the rankings are aggregated (top). For aggregate-then-rank methods, spectra are first aggregated to form a consensus spectrum, and each such spectrum is then ranked against the candidates (bottom).

#### S3.4 Comparative study between FormSpec and BinnedSpec

We compare the performance between using FormSpec, the MVP spectra encoder, and BinnedSpec, a common spectra encoder architecture used in prior works[S1,S8] (Table S3 and Figure S6). Further comparisons on the spectra-candidate similarity and ranking variability within spectra of the same molecule are shown in Figure S7.

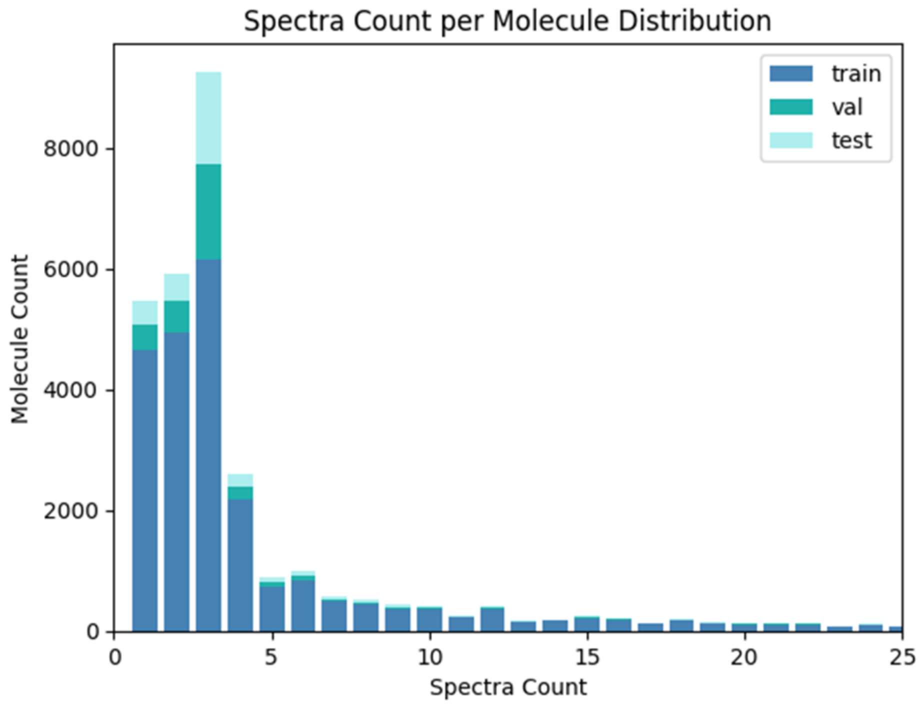


Figure S4: Distribution of spectra count per molecule across train, validation, and test split.

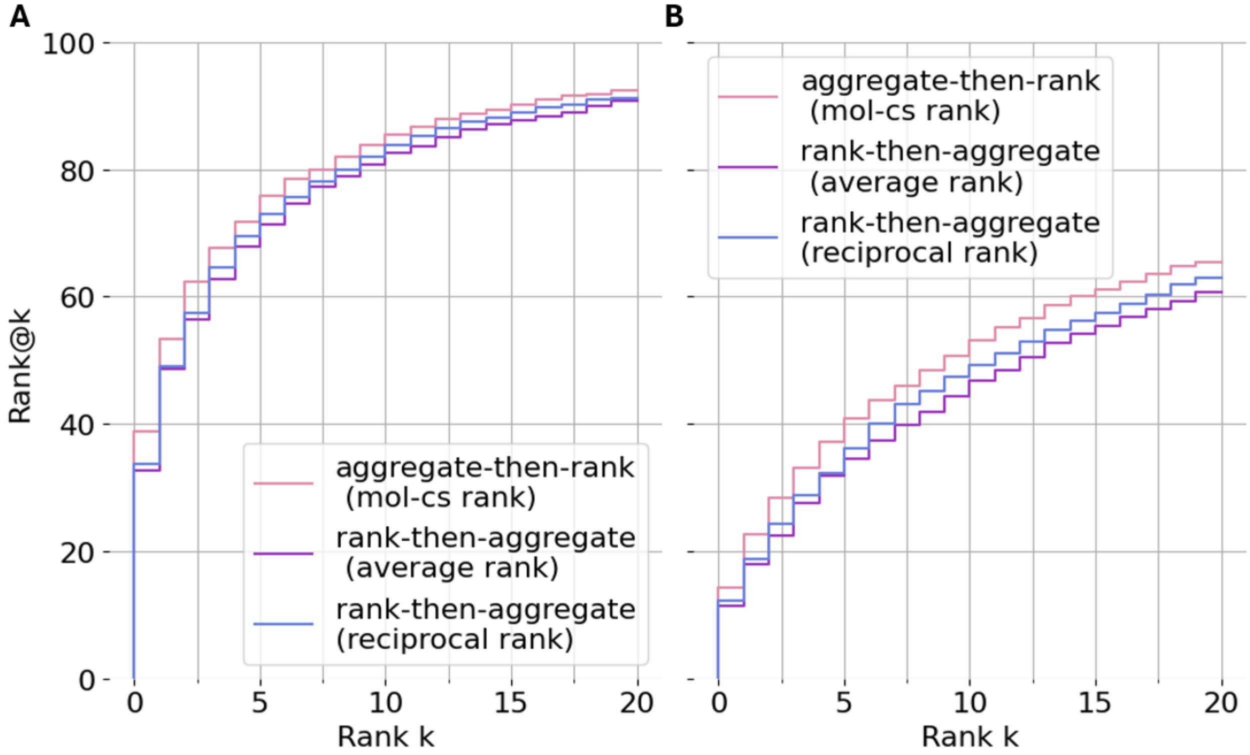


Figure S5: Staircase plots of ranking results showing that aggregate-then-rank outperforms rank-then-aggregate methods when retrieving candidates by mass (A) and formula (B).

Table S3: Comparative study on the spectra features. The best performance is **highlighted** and the best performance based on the mol-s ranking views is underlined.

| Ranking views | Spectra feature | A. MassSpecGym rank@1 (↑) | rank@5 (↑) | rank@20 (↑) | B. [M+H]+ subset rank@1 (↑) | rank@5 (↑) | rank@20 (↑) |
|---|---|---|---|---|---|---|---|
| I. Candidates by Mass | | | | | | | |
| mol-s | FormSpec | 26.37 | 58.90 | 86.88 | 29.73 | 62.18 | 88.3 |
| | BinnedSpec | 12.29 | 29.39 | 51.03 | 12.38 | 30.31 | 52.82 |
| mol-cs | FormSpec | **35.99** | **67.97** | **91.92** | **36.93** | **72.01** | **92.87** |
| | BinnedSpec | 16.63 | 40.07 | 65.26 | 16.81 | 39.33 | 64.51 |
| II. Candidates by Formula | | | | | | | |
| mol-s | FormSpec | 11.10 | 31.14 | 61.99 | 12.27 | 33.2 | 61.85 |
| | BinnedSpec | 11.48 | 29.91 | 57.78 | 10.03 | 27.61 | 55.89 |
| mol-cs | FormSpec | 13.96 | **36.88** | **68.12** | **14.65** | **38.99** | **68.31** |
| | BinnedSpec | **15.05** | 35.48 | 64.17 | 12.83 | 31.43 | 63.46 |

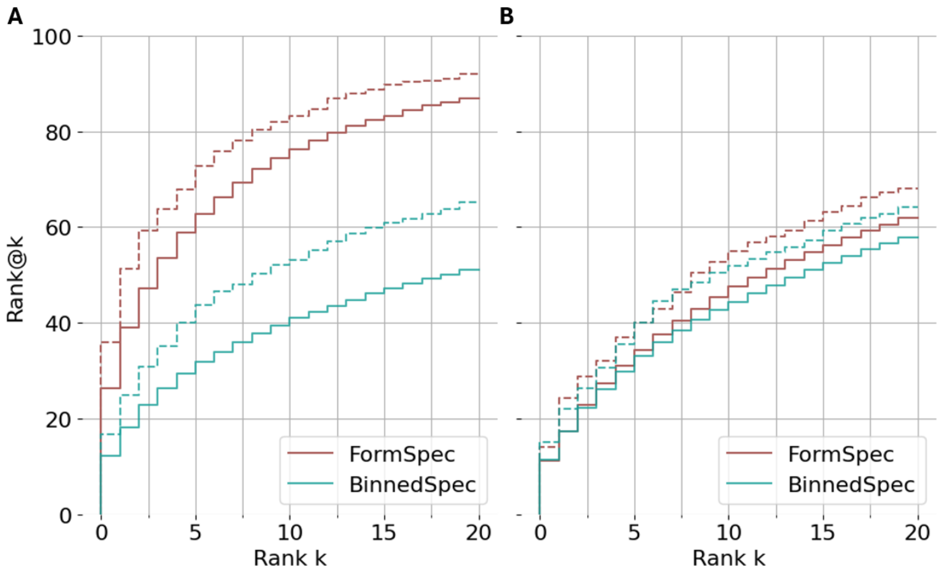


Figure S6: Staircase plots of ranking performance using BinnedSpec vs FormSpec for candidates by mass(A) and formula(B) with solid lines for mol-s ranking views and dashed lines for mol-cs ranking views.

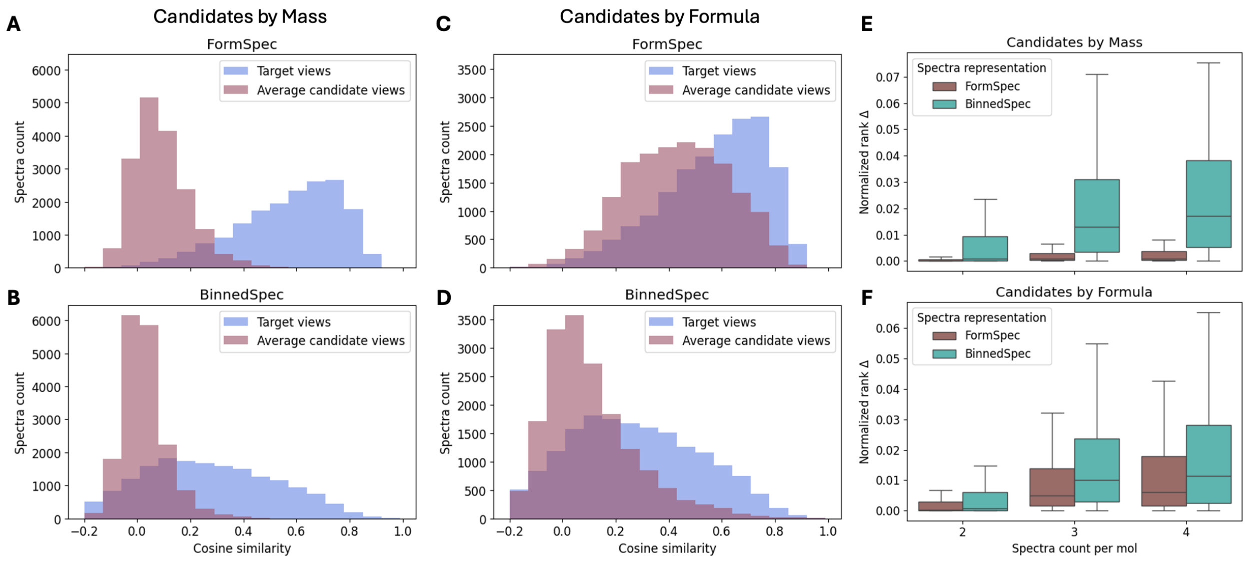


Figure S7: Comparative study on spectra representations. Histograms of spectra-target molecule cosine similarity vs average spectra-candidate cosine similarity when using FormSpec and BinnedSpec on candidates by mass (A,B) and formula (C,D). Boxplots of rank variability using FormSpec vs BinnedSpec for test molecules with varying test spectra count when retrieving candidates by mass (E) and formula (F).

#### S3.5 Ablation study on training views

We conduct an ablation study to show the contribution of each view used in training (Table S4 and Figure S8).

#### S3.6 Case study

We include additional figures in support of using MVP to compare cultured and conventional meat metabolic profiles.[S9] We use MVP to rank the candidates, and the statistics of the annotation scores are shown in Figure S9A and S9B. The abundance across pathways based on the features identified by the original study is shown in Figure S9C.

Table S4: Ablation study on the training views. The best performance is **highligted**, and the best performance based on the mol-s ranking views is underlined.

| Ranking views | Training views | A. MassSpecGym rank@1 (↑) | A. MassSpecGym rank@5 (↑) | A. MassSpecGym rank@20 (↑) | B. [M+H]+ subset rank@1 (↑) | B. [M+H]+ subset rank@5 (↑) | B. [M+H]+ subset rank@20 (↑) |
|---|---|---|---|---|---|---|---|
| I. Candidates by Mass | | | | | | | |
| mol-s | mol-fp-s-cs | 26.37 | 58.90 | 86.88 | 29.73 | 62.18 | 88.30 |
| | mol-s-cs | 25.60 | 59.13 | 86.91 | 25.56 | 59.49 | 87.42 |
| | mol-fp-s | 26.48 | 58.85 | 86.49 | 29.22 | 61.97 | 88.75 |
| | mol-s | 25.46 | 59.01 | 86.69 | 27.83 | 59.33 | 86.84 |
| mol-cs | mol-fp-s-cs | **35.99** | 67.97 | **91.92** | **36.93** | **72.01** | **92.87** |
| | mol-s-cs | 33.17 | **69.26** | 90.66 | 30.14 | 68.79 | 91.24 |
| II. Candidates by Formula | | | | | | | |
| mol-s | mol-fp-s-cs | 11.10 | 31.14 | 61.99 | 12.27 | 33.20 | 61.85 |
| | mol-s-cs | 10.53 | 29.92 | 59.83 | 10.03 | 28.63 | 58.80 |
| | mol-fp-s | 11.34 | 31.33 | 61.20 | 11.78 | 32.77 | 61.74 |
| | mol-s | 10.79 | 30.18 | 59.23 | 10.65 | 29.14 | 58.09 |
| mol-cs | mol-fp-s-cs | **13.96** | **36.88** | **68.12** | **14.65** | **38.99** | **68.31** |
| | mol-s-cs | 13.20 | 36.85 | 66.46 | 11.83 | 35.25 | 64.11 |

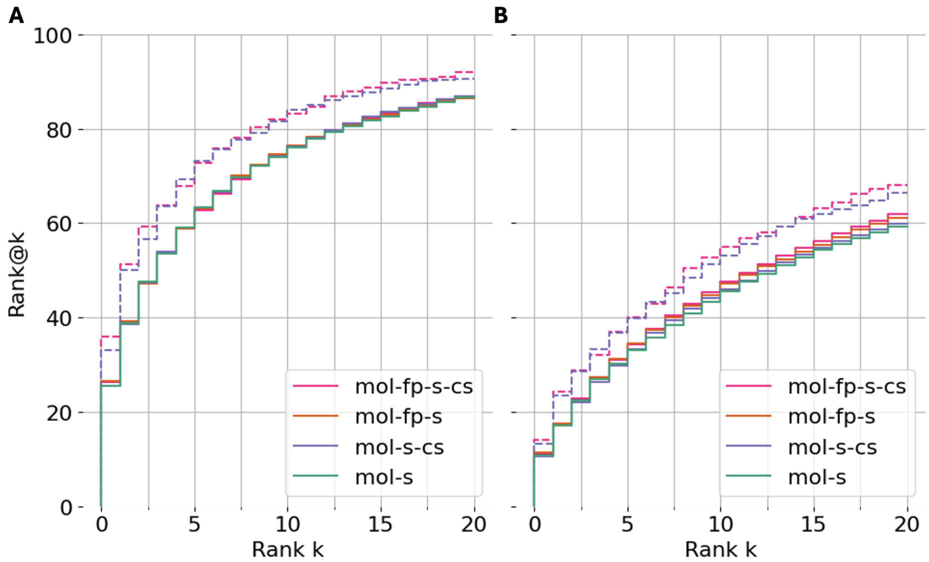


Figure S8: Staircase plots of ranking performance when trained on various views and evaluated on candidates by mass (A) and formula (B) with solid lines for mol-s ranking views and dashed lines for mol-cs ranking views.

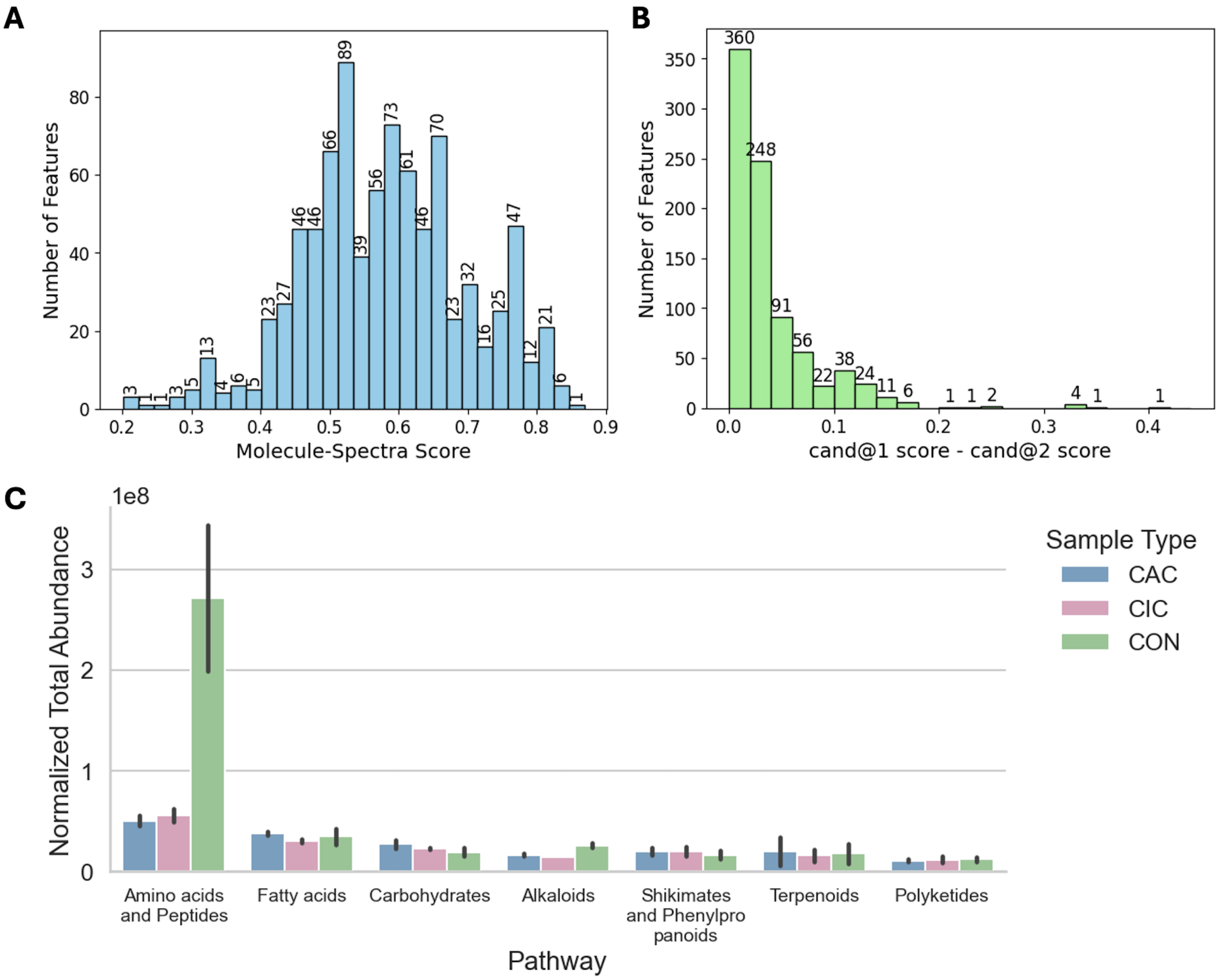


Figure S9: Case study results. A) Distribution of molecule-spectra scores of the top-ranked molecule identified by MVP. B) Distribution of score differences between the top- and second-ranked candidates. C) Total abundance of each pathway per sample type based on the annotations from the original work.
